# Presynaptic mitochondria calcium uniporter promotes auditory temporal processing during sustained high-rate activity

**DOI:** 10.64898/2026.06.21.733581

**Authors:** Guanyu Li, Ruili Xie

**Author notes:** **Corresponding author**: Ruili Xie.

## Abstract

Mitochondrial calcium uniporter (MCU) uptakes calcium into mitochondria to maintain intracellular calcium homeostasis, malfunction of which has been implicated in altered neuronal signaling and disease. Its role in synaptic transmission remains understudied, especially in intact neural circuits. We investigated MCU function at the auditory nerve endbulb of Held synapse and postsynaptic bushy neurons in the cochlear nucleus, using age-matched control and MCU knockout (KO) mice of either sex. Whole-cell voltage-and current-clamp recordings were acquired from acute brain slices to examine synaptic transmission and postsynaptic responses. We found that basal synaptic properties at the endbulb of Held were unchanged in MCU KO mice, whereas synaptic transmission during sustained high-rate activity was significantly altered with a shift toward increased asynchronous release. Similarly, MCU deficiency did not change the intrinsic membrane properties of postsynaptic bushy neurons, but significantly reduced the temporal precision of auditory nerve evoked spikes trains at high rates. These results demonstrate that MCU is largely dispensable under low-rate activity, presumably because its activation requires relatively high calcium concentrations. In contrast, during sustained high-rate activity, MCU becomes an important regulator of synaptic function by reducing asynchronous neurotransmitter release under elevated intracellular calcium. Particularly in the auditory system, where neurons routinely fire at high rates, MCU promotes temporal processing and thereby plays a key role in supporting auditory function. It suggests that impaired MCU function under pathological conditions may be an important mechanism underlying central auditory processing deficits, and consequently contributes to hearing loss.

**SIGNIFICANT STATEMENT:** MCU uptakes calcium into mitochondria to regulate intracellular calcium, yet its contribution to synaptic transmission and neural processing remain understudied. Using a MCU KO mouse model, we investigated the role of MCU in synaptic transmission and postsynaptic responses at the endbulb of Held synapses and postsynaptic bushy neurons in the cochlear nucleus. The results showed that MCU plays little role in basal synaptic transmission, but significantly reduces asynchronous vesicle release during sustained high-rate activity, consequently improving temporal precision of the signal processing. These findings define the roles of MCU under different activity levels at the intact endulb/bushy connection, and suggest that impaired MCU function may be a key mechanism underlying central auditory processing deficits, and consequently contributes to hearing loss.

## INTRODUCTION

Mitochondria calcium uniporter (MCU) is a calcium-selective ion channel located on the inner membrane of mitochondria (Alevriadou et al., 2021). It mediates calcium uptake into mitochondria and helps maintain intracellular calcium homeostasis, dysfunction of which leads to pathological diseases including cancer (Vultur et al., 2018), neurodegenerative disorder (Liao et al., 2017; Ashleigh et al., 2023), and metabolic diseases (Li et al., 2023). In synaptic transmission, the fundamental process underlying neural communication, MCU has been reported to regulate synaptic plasticity in various brain circuits, including via postsynaptic mechanisms in hippocampal CA2 neurons (Pannoni et al., 2025) and spinal dorsal horn neurons (Kim et al., 2011), and via presynaptic mechanism in hippocampal mossy fibre synapses (Devine et al., 2022). Effective calcium clearance is critical at the presynapse (Devine and Kittler, 2018) for maintaining synaptic transmission, especially during sustained high-rate activity, when repetitive calcium influx leads to substantial calcium accumulation. MCU-mediated calcium uptake accounts for up to 40% of presynaptic calcium clearance under these conditions (Kim et al., 2005; Shutov et al., 2013). However, the roles of MCU in synaptic transmission remain understudied, especially in intact neural circuits.

The cochlear nucleus (CN) constitutes the first relay station of the central auditory pathway, where auditory nerve (AN) inputs are first processed within the brainstem, including the giant axosomatic synapse known as the endbulb of Held innervating CN principal bushy neurons (Ryugo and Fekete, 1982; Manis et al., 2011). These synapses have long served as a canonical model for studying synaptic transmission and are capable of sustaining exceptionally high firing rates at physiologically relevant sound levels (Kiang et al., 1965; Taberner and Liberman, 2005; Wen et al., 2009). These synaptic terminals contain a high density of mitochondria (Ryugo et al., 1996; Ryugo et al., 1997), reflecting their high energetic and calcium-buffering demands, which makes them an ideal model for investigating the role of MCU in synaptic function. Furthermore, the endbulb of Held / bushy neuron connection is specialized in processing fine temporal information of sound critical for auditory function (Joris et al., 1994b; Shannon et al., 1995; Joris and Yin, 2007; Walton, 2010). A careful investigation of MCU function at this neural connection is essential for understanding its role in central auditory signal processing and its broader contributions to brain function.

In this study, we used control and MCU knockout (KO) mice to address two primary questions: (1) how does MCU contribute to synaptic transmission at the endbulb of Held, and (2) how does MCU deficiency affect signal processing in postsynaptic bushy neurons, potentially contributing to central auditory processing deficits and subsequently hearing loss. We found that MCU deficiency does not change the basal synaptic properties of endbulb of Held, or intrinsic membrane properties of postsynaptic bushy neurons. During sustained high-rate activity, synaptic transmission at the endbulb of Held in MCU KO mice showed similar total vesicle release but a proportionally greater asynchronous release as activity progressed. Further evaluation of the spike output of the bushy neurons revealed that while the firing rate of AN evoked spikes during prolonged activity remained unchanged in MCU KO mice, their temporal precision was significantly compromised. These results demonstrate that MCU is largely dispensable for basal synaptic transmission, presumably because its activation requires relatively high calcium concentrations. In contrast, during sustained high-rate activity, MCU becomes an important regulator of synaptic signal processing by shaping neurotransmitter release under elevated intracellular calcium. Particularly in the auditory system, where neurons routinely fire at high rates, MCU promotes temporal processing and thereby plays a key role in supporting auditory function. It suggests that impaired MCU function under pathological conditions may be an important mechanism underlying central auditory processing deficits, and consequently contributes to hearing loss.

## MATERIALS AND METHODS

### Research animals

MCU heterozygous (MCU^+/-^) mice ON FVB/NJ background were obtained courtesy of Dr. Ruben Stepanyan from Case Western Reserve University (Manikandan et al., 2021), housed and bred at the animal facility at The Ohio State University. Homozygous MCU KO (MCU^-/-^) mice were produced and used at 3-4 months of age in this study to investigate the effect of MCU deficiency on synaptic transmission and signal processing. Due to close genomic location, MCU KO allele is tightly linked to *Cdh23^ahl^* allele, which is known to lead to early onset age-related hearing loss. To minimize this potential confounding effect, we limited the age of the experimental mice to no more than 4 months, before significant hearing loss develops due to *Cdh23^ahl^*allele. Age-matched littermate MCU heterozygous mice were used as control, which share a genetic background more similar to the MCU KO mice (Manikandan et al., 2021). These control mice retain reduced yet robust MCU mRNA expression (hence MCU function), develop normally, and exhibit relatively normal hearing. Mice of either sex were used. All experiments were conducted under the guidelines of the protocol #2018A00000055-R2 approved by the Institutional Animal Care and Use Committee of The Ohio State University.

### Auditory brainstem response (ABR)

Hearing status of the mice was evaluated by measuring ABR to clicks as described previously (Wang et al., 2021). Mice were deeply anesthetized using a cocktail of ketamine (100 mg/kg) and xylazine (10 mg/kg) via intraperitoneal injection, and placed on a thermo-controlled heating pad inside a custom-made sound-attenuating chamber to maintain body temperature at 36 *^◦^*C. ABR to clicks were recorded using an RZ6-A-P1 system with BioSigRz software (Tucker-Davis Technologies). Clicks (0.1 ms, monophasic with alternating phase) were delivered at 21 times/s through a free-field MF1 magnetic speaker (Tucker-Davis Technologies) located 10 cm away from the pinna. Two recording needle electrodes were respectively placed at the ipsilateral pinna and vertex, whereas a third ground electrode was placed at the rump. ABR at each sound level was recorded 512 times and averaged. ABR threshold was determined as the lowest sound level that evoked recognizable ABR waveforms.

### Brain slice preparation

Acute brain slices were prepared as previously described (Xie, 2016; Xie and Manis, 2017). Artificial cerebrospinal fluid (ACSF) was freshly made with the following recipe (in mM): 122 NaCl, 3.0 KCl, 1.8 CaCl_2_, 1.5 MgSO_4_, 1.25 KH_2_PO_4_, 20 glucose, 25 NaHCO_3_, 0.4 ascorbic acid, 2 sodium pyruvate, and 3 myoinositol. The ACSF solution was pre-warmed to 34 °C in water bath and gassed with 95% O_2_ and 5% CO_2_. Under deep anesthesia, mice were decapitated to retrieve brainstem. Parasagittal slices containing the CN were cut at the thickness of 250 µm using a vibratome slicer (model VT1200S Microtome, Leica Biosystems), and were incubated in ACSF solution at 34 °C for 30 to 40 minutes to recover.

### Electrophysiological recording

After incubation, individual brain slice was transferred to a recording chamber under a microscope (Axio Examiner, Carl Zeiss), bathed in ACSF with a flowing rate of 2-3 ml/min. Whole-cell recording under either voltage clamp mode or current clamp mode was performed from bushy neurons in the middle and high frequency regions of the anteroventral CN (AVCN). Recording electrodes were pulled using a P-2000 micropipette puller (Sutter Instrument). For voltage clamp recording, electrode pipette was filled with Cs^+^-based internal solution containing (in mM): 105 CsMetSO_3_, 35 CsCl, 5 EGTA, 10 HEPES, 4 MgATP, 0.3 GTP, 10 phosphocreatine, and 3 QX-314 chloride. For current clamp recording, electrode pipette was filled with K^+^-based internal solution containing (in mM): 126 potassium gluconate, 6 KCl, 2 NaCl, 10 HEPES, 0.2 EGTA, 4 MgATP, 0.3 GTP and 10 Tris-phosphocreatine. Both internal solutions had an osmolarity of 296 mOsm/L, with PH adjusted to 7.20. Alexa-Fluor 594 (0.01% by weight) was added to the internal solution to fill the target neurons to allow visualization of cellular morphology upon completion. Two µM strychnine was added to the bath ACSF to block glycinergic inhibitory synaptic transmission. Electrophysiological data were acquired using a Multiclamp 700B amplifier, a Digidata 1550B Acquisition System, and pClamp 11 software (Molecular Devices). To drive synaptic transmission, AN was stimulated by a 0.1 ms voltage pulse via a 75-µm-diameter concentric stimulating electrode (Frederick Haer Company). To ensure maximum stimulation, the stimulus intensity was set at 20% to 50% above the voltage level needed to trigger maximum evoked EPSCs (eEPSCs) or reliable spikes. For voltage clamp recording, cells were held at-67mV with series resistance compensated by around 75%. Spontaneous EPSC (sEPSC) events were recorded first, followed by eEPSCs triggered by paired-pulse stimulation at the AN to study the synaptic transmission under quiescence. Subsequently, synaptic properties under low-and high-rate activities were studied using 50-pulse stimulus trains at 100 and 400 Hz, followed by a single post-train stimulation at various delays between 10 and 1500 ms after the end of each train to evaluate synaptic recovery. For current clamp recording, data acquisition started with direct current step injection evoked responses to assess the intrinsic membrane properties of the target bushy neurons. Then, responses to short (50 pulses) or long (900 pulses) AN stimulus trains at both 100 Hz and 400 Hz were acquired to investigate AN-evoked firing properties in target bushy neurons. All recordings were made at 34 *^◦^*C. A junction potential of-7 mV for voltage-clamp recording, or-12 mV for current clamp recording was corrected in all reported voltages.

### Data analysis

sEPSC events were analyzed using Clampfit software (Molecular Devices). All other electrophysiological data were analyzed using custom-written programs in either MATLAB (MathWorks) or Igor Pro (WaveMetrics) as described in previous work (Xie and Manis, 2013; Xie, 2016; Xie and Manis, 2017), and specified in the Results section.

### Statistical analysis

All statistical analyses were performed using Prism (GraphPad Software). Unpaired Student’s t test (or Mann–Whitney test when the data population failed to pass the normality test), and two-way ANOVA were used as stated. Data are presented as mean ± standard deviation (SD) throughout the manuscript, except certain figures where mean ± standard error of the mean (SEM) were used for visual clarity as stated.

## RESULTS

### MCU KO mice show severe hearing loss and altered neural processing in the cochlear nucleus

MCU is essential for hearing and its deficiency contributes to early onset hearing loss (Manikandan et al., 2021). We evaluated hearing function in 3-month-old MCU KO mice by recording their ABRs to clicks at intensities from 15 to 90 dB SPL. Compared to age-matched control littermates, MCU KO mice showed significantly elevated ABR threshold and reduced Wave I and Wave II amplitudes in response to clicks (Fig. 1A). On average, the click threshold was 35 *±* 4 dB SPL in control mice (n=12) and was significantly higher at 64 *±* 12 dB SPL in MCU KO mice (n=18) (Fig. 1B; Welch’s t test: P *<* 0.0001). ABR wave I amplitude was greatly reduced in MCU KO mice at sound levels above 40 dB, whereas no significant difference was observed at lower intensities (Fig. 1C, two-way ANNOVA, sound intensity effect: F_(15,448)_ = 69.26, P *<* 0.0001; mice strain effect: F_(1, 448)_ = 692.6, P *<* 0.0001; interaction: F_(15, 448)_ = 30.58, P *<* 0.0001; Sidak’s multiple comparisons test: P *<* 0.0001 for all levels above 45 dB SPL, P = 0.0014 at 40 dB SPL, P *>* 0.5 for all level below 40 dB SPL). ABR wave II amplitude was also reduced in a similar pattern, with significantly smaller response in MCU KO mice at 45 dB SPL and above, and no difference at lower intensities (Fig. 1D; two-way ANNOVA, sound intensity effect: F_(15, 448)_ = 24.53, P *<* 0.0001; mice strain effect: F_(1, 448)_ = 259.0, P *<* 0.0001; interaction: F_(15, 448)_ = 10.80, P *<* 0.0001; Sidak’s multiple comparisons test: P *<* 0.0001 at all levels above 45 dB, P *>* 0.5 for sound levels at 40 dB SPL and below). These results demonstrate that MCU KO mice exhibit severe hearing impairment, confirming previous report (Manikandan et al, 2021). It is worth noting that the amplitude of wave I increased monotonically with sound intensity in both control and MCU KO mice (Fig. 1C), whereas the amplitude of wave II increased non-monotonically in control but monotonically in KO mice. Since ABR wave II primarily reflects neural activity in the CN (Melcher and Kiang, 1996; Wang et al., 2025), the changed dynamics in ABR wave II amplitude across sound intensity levels between control and MCU KO mice suggest that MCU deficiency associates with altered neural processing within the CN.

**Figure 1:**
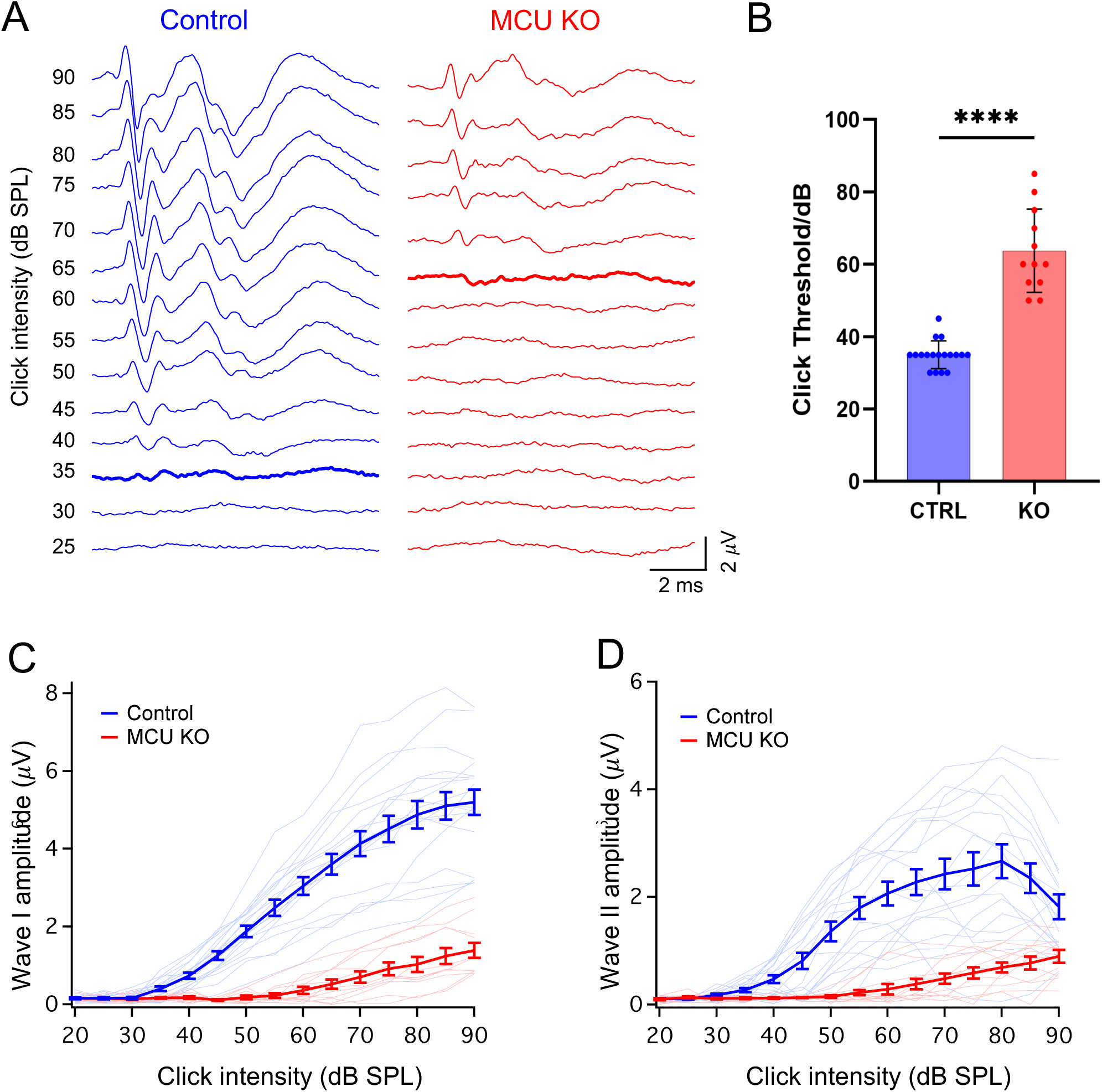
MCU KO mice exhibit severe hearing impairment at 3 months. **A**. Example ABR waveforms from control and MCU KO mice in response to clicks. Thick traces mark responses at hearing threshold. **B**. Comparison of click threshold between control and MCU KO mice. ****P *<* 0.0001. **C-D**. Growth curves of the ABR wave I amplitude (**C**) and wave II amplitude (**D**) across sound intensity. Thin lines represent individual mice; thick lines represent mean ± SEM.

### MCU deficiency does not affect basal synaptic transmission at the endbulb of Held synapse

The CN receives all sound information from the AN through glutamatergic synapses, including the endbulb of Held, a prominent and specialized synapse that plays a crucial role in processing fine temporal information essential for auditory tasks (Manis et al., 2011). To investigate the role of MCU in neural processing, we studied the basal synaptic transmission at the endbulb of Held in control and MCU KO mice. Whole-cell voltage clamp recording was acquired from postsynaptic bushy neurons to evaluate synaptic properties including spontaneous EPSCs (sEPSCs) under quiescence, single and paired-pulse evoked EPSCs (eEPSCs). Cs-based internal solution with QX-314 was used to block voltage-gated potassium and sodium channels, respectively. As shown in Fig. 2A, spontaneous synaptic transmission in control and MCU KO mice showed similar sEPSC release rate and sEPSC amplitude. On average, sEPSC frequency was 3.7 ± 1.4 Hz (n = 22) in control and 4.7 ± 3.2 Hz (n = 26) in MCU KO mice (Fig. 2B; Mann-Whitney test: P = 0.8819); and the average sEPSC amplitude was-112.9 ± 49.8 pA in control and-134.3 ± 59.0 pA in MCU KO mice (Fig. 2C; Mann-Whitney test: P = 0.0840). sEPSC decay time constant was 0.19 ± 0.06 ms (n = 22) in control and 0.18 ± 0.04 ms (n = 26) in MCU KO mice (Fig. 2D; Welch’s t test, P = 0.5369). These results indicate that MCU deficiency does not alter basal synaptic properties at the endbulb of Held, including the quantal size of synaptic vesicles and kinetics of postsynaptic glutamatergic receptors.

**Figure 2:**
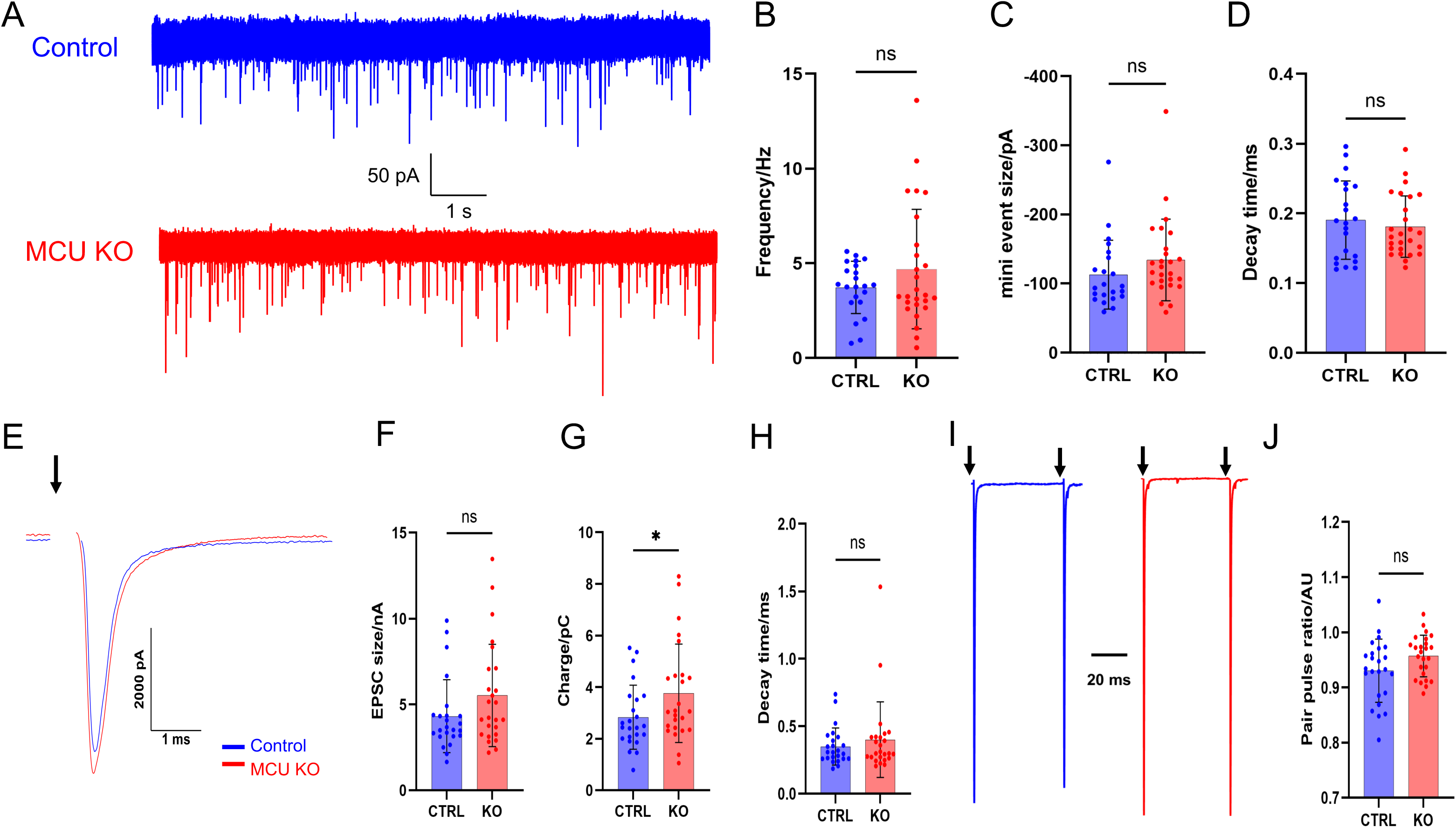
MCU deficiency does not affect basal synaptic transmission. **A**: Example sEPSC events in bushy neurons from control and MCU KO mice. **B-D**: Summary of sEPSC event frequency (**B**), average peak amplitude (**C**), and decay time constant (**D**). ns, not significant. **E**: Example eEPSCs to single AN stimulation from control and MCU KO mice. Arrow marks the stimulus onset. **F-H**: Summary of eEPSC peak amplitude, charge transfer, and decay time constant. ns, not significant; *P *<* 0.5. **I**: Example eEPSCs to paired-pulse stimulation with a 50 ms inter-pulse interval. **J**: Summary paired-pulse ratio. ns, not significant.

We then studied evoked synaptic response by stimulating the AN with single or paired voltage pulses to trigger eEPSCs (Fig. 2E-J). There was a trend that the peak amplitude of single eEPSCs was larger in MCU KO mice, suggesting that quantal content may be increased (Fig. 2E). However, the difference was not statistically significant (Fig. 2F; Control, 4.3 ± 2.1 nA, n = 24; MCU KO, 5.5 ± 3.0 nA, n = 25; Mann-Whitney test, P = 0.1412). We also measured the total charge transfer of the single eEPSC by integrating the area under the curve of the EPSC within a 5 ms window after the rising onset of the eEPSC. On average, eEPSCs from MCU KO synapses showed significantly larger total charge release than those from control synapses (Fig. 2G; Control, 2.83 ± 1.24 pC, n = 24; MCU KO, 3.76 ± 1.91 pC, n = 25; Welch’s t test: P = 0.0487). There was no difference in eEPSC decay time constant between two groups (Fig. 2H; Control, 0.35 ± 0.14 ms, n = 24; MCU KO, 0.40 ± 0.28 ms, n = 25; Mann-Whitney test, P = 0.7248). For paired-pulse eEPSCs with a 50 ms inter-stimulus interval, we calculated the paired pulse ratio and found no significant difference between control and MCU KO mice (Fig. 2I-J; Control, 0.93 *±* 0.06, n = 24; MCU KO, 0.96 *±* 0.04, n = 25; Welch’s t test: P = 0.0637). Overall, we conclude that MCU deficiency does not alter basal synaptic transmission at the endbulb of Held synapses, including quantal content and release probability of synaptic vesicles under quiescence.

### MCU deficiency shifts synaptic transmission toward increased asynchronous release during sustained high-rate activity

MCU regulates intracellular calcium by uptaking calcium into mitochondria under high calcium level, which occurs at synaptic terminals during high-rate activity. AN is highly active and can fire up to 400 Hz under physiological conditions over prolonged time (Kiang et al., 1965; Taberner and Liberman, 2005; Wen et al., 2009). Synaptic transmission during such high-rate activity requires repetitive calcium influx and efficient calcium clearance, including MCU-mediated calcium uptake into mitochondria. To further study MCU function, we investigated synaptic transmission at the endbulb of Held synapse in response to 50-pulse stimulus trains at 100 and 400 Hz in control and MCU KO mice. At 100 Hz, AN stimulation evoked robust eEPSCs at the beginning of the train in both groups, followed by eEPSCs with reduced amplitude due to synaptic depression (Fig. 3A). However, synaptic depression was greater in MCU KO mice during the second half of the trains (Fig. 3C). We calculated the steady state depression by averaging the amplitude of the last 25 eEPSCs relative to the first eEPSC of the train. On average, steady-state depression was 0.65 *±*0.15 (n = 21) in control and 0.55 *±* 0.14 (n = 26) in MCU KO mice (Fig. 3C; Welch’s t test, P = 0.0293). Expectedly, eEPSCs depressed more in 400 Hz train than 100 Hz, and the depression was most profound in MCU KO mice (Fig. 3B). The steady-state depression was on average 0.33 *±* 0.13 (n = 20) in control and 0.18 *±* 0.09 (n = 26) in MCU KO mice (Fig. 3D; Welch’s t test, P *<* 0.0001).

**Figure 3:**
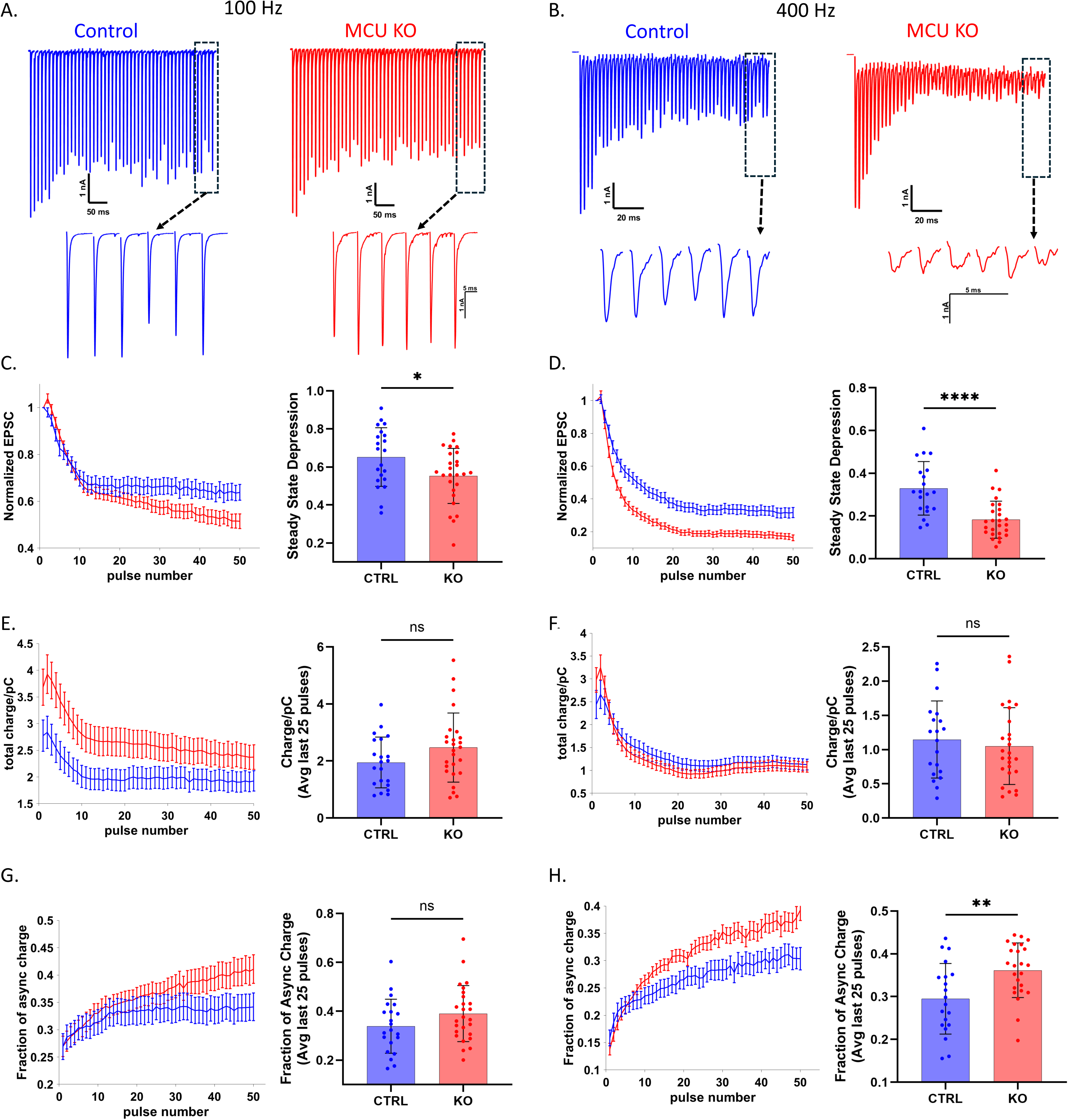
MCU deficiency impairs synaptic transmission during sustained high-rate activity. **A-B**: Representative eEPSCs to AN stimulus trains at 100 Hz (**A**) and 400 Hz (**B**) from control and MCU KO mice. **C-D**: Normalized eEPSC amplitude throughout the stimulus trains and summary of steady-state depression at 100 Hz (**C**) and 400 Hz (**D**). *P*<*0.05, ****P*<*0.0001. **E-F**: eEPSC charge transfer throughout the stimulus trains and mean charge transfer of the last 25 eEPSCs of the trains at 100 Hz (**E**) and 400 Hz (**F**). ns, not significant. **G-H**: Fraction of asynchronous release within each eEPSC throughout the stimulus trains and mean asynchronous fraction during the last 25 eEPSCs of the trains at 100 Hz (**G**) and 400 Hz (**H**). ns, not significant. **P*<*0.01.

The peak amplitude of eEPSC reflects the synchronous component of the synaptic response, but does not take account of the synaptic response from asynchronous vesicle release, which is likely more affected by calcium accumulation (Kaeser and Regehr, 2014), hence MCU function. Therefore, we further calculated the charge transfer of each eEPSC to quantify the total vesicle release throughout the trains (Fig. 3E, F). Since the charge transfer of single eEPSC differ between control and MCU KO mice (Fig. 2G), the charge transfer data in trains were not normalized. Consistently, eEPSC charge transfer was larger at the beginning of the trains and decreased in size toward the end of the trains at both 100 and 400 Hz (Fig. 3E-F). Unexpectedly, we found that the average charge transfer of the last 25 eEPSCs was not significantly different between control and MCU KO mice at both 100 Hz (Fig. 3E; Control, 1.95 ± 0.89 pC, n = 21; MCU KO, 2.47 *±* 1.21 pC, n = 26; Welch’s t test, P = 0.0954) and 400 Hz (Fig. 3F; Control, 1.15 ± 0.56 pC, n = 20; MCU KO, 1.05 ± 0.56 pC, n = 26; Welch’s t test, P = 0.5714). To further characterize asynchronous release within each eEPSC, we defined asynchronous release as the charge transfer outside of a 1 ms window surrounding the eEPSC peak (0.3 ms before and 0.7 ms after the peak), and calculated the fraction of asynchronous release of all individual eEPSCs throughout the trains. The fraction of asynchronous release was similar between control and MCU KO synapses at the beginning of the trains, but progressively became higher toward the later stage of the trains in MCU KO mice at both 100 and 400 Hz (Fig. 3G, H). During the last 25 pulses, the average fraction of asynchronous release was not significantly different between two groups but trended higher in MCU KO at 100 Hz (Fig. 3G; Control, 33.87% *±* 11.96%, n = 21; MCU KO, 39.04% *±* 11.47%, n = 26; Welch’s t test, P = 0.1244). At 400 Hz, asychronous release fraction was significantly higher in MCU KO synapses (Fig. 3H; Control, 29.49% *±* 8.25%, n = 20; MCU KO, 36.16% *±*6.38%, n = 24; Welch’s t test, P = 0.0055). Collectively, these results suggest that MCU deficiency shifts synaptic transmission toward higher asynchronous release during sustained high-rate activity, presumably due to enhanced calcium accumulation at the synaptic terminal. The total synaptic release during the steady state of the trains, however, remained largely unchanged (Fig. 3E-F).

We calculated the readily releasable pool (RRP) size using 400 Hz eEPSC trains based on methods from Schneggenburger et al. (1999) and Elmqvist and Quastel (1965) (Fig. 4). First, the number of vesicles being released was calculated as eEPSC peak amplitude divided by the average sEPSC amplitude. For the method by Schneggenburger et al. (1999), the portion of the cumulative plot at the steady-state phase (15th to 50th eEPSC) was fitted into a line and back-extrapolated to time 0 to estimate the RRP size (Fig. 4A). No significant difference was found in RRP size at the endbulb of Held between control and MCU KO mice (Fig. 4B; Control, 173 *±*100, n = 17; MCU KO, 182 *±*84, n = 25; Mann-Whitney test, P = 0.4616). For the method by Elmqvist and Quastel (1965), individual eEPSC releases across the train were plotted against cumulative releases, and the initial linear phase (first 4-5 releases) was used for linear fitting. The RRP size was estimated from the x-axis intercept of the fitted line (Fig. 4C), which was also similar between control and MCU KO mice (Fig. 4D; Control, 388 *±* 244, n = 16; MCU KO, 310 *±* 192, n = 25; Mann-Whitney test, P = 0.3073). Second, we calculated the RRP size based on the charge transfer of each eEPSC throughout the trains in order to additionally take account of the asynchronous response component. The number of vesicles released within each eEPSC was estimated by dividing the total charge of each eEPSC by the mean charge of the average sEPSC. Based on the method by Schneggenburger et al. (1999), the estimated RRP size was 347 ± 188 (n = 17) in control and 339 ± 116 (n = 25) in MCU KO (Fig. 4E-F; Mann-Whitney test, P = 0.8592). Consistently, the estimated RRP size using Elmqvist and Quastel (1965) method was 1214 ± 797 (n = 14) in control and 1082 ± 650 (n = 25) in MCU KO (Fig. 4G-H; unpaired t test, P = 0.6022). Three cells from the control mice were excluded from the analysis due to the non-linear pattern of their initial cumulative plot, which was against the assumption of initial linear RRP release of the method and resulted negative RRP size estimates. These results showed that the initial RRP size was not changed in MCU KO mice, further supporting the findings in Fig. 2 that MCU deficiency does not alter the basal synaptic properties of endbulb of Held.

**Figure 4:**
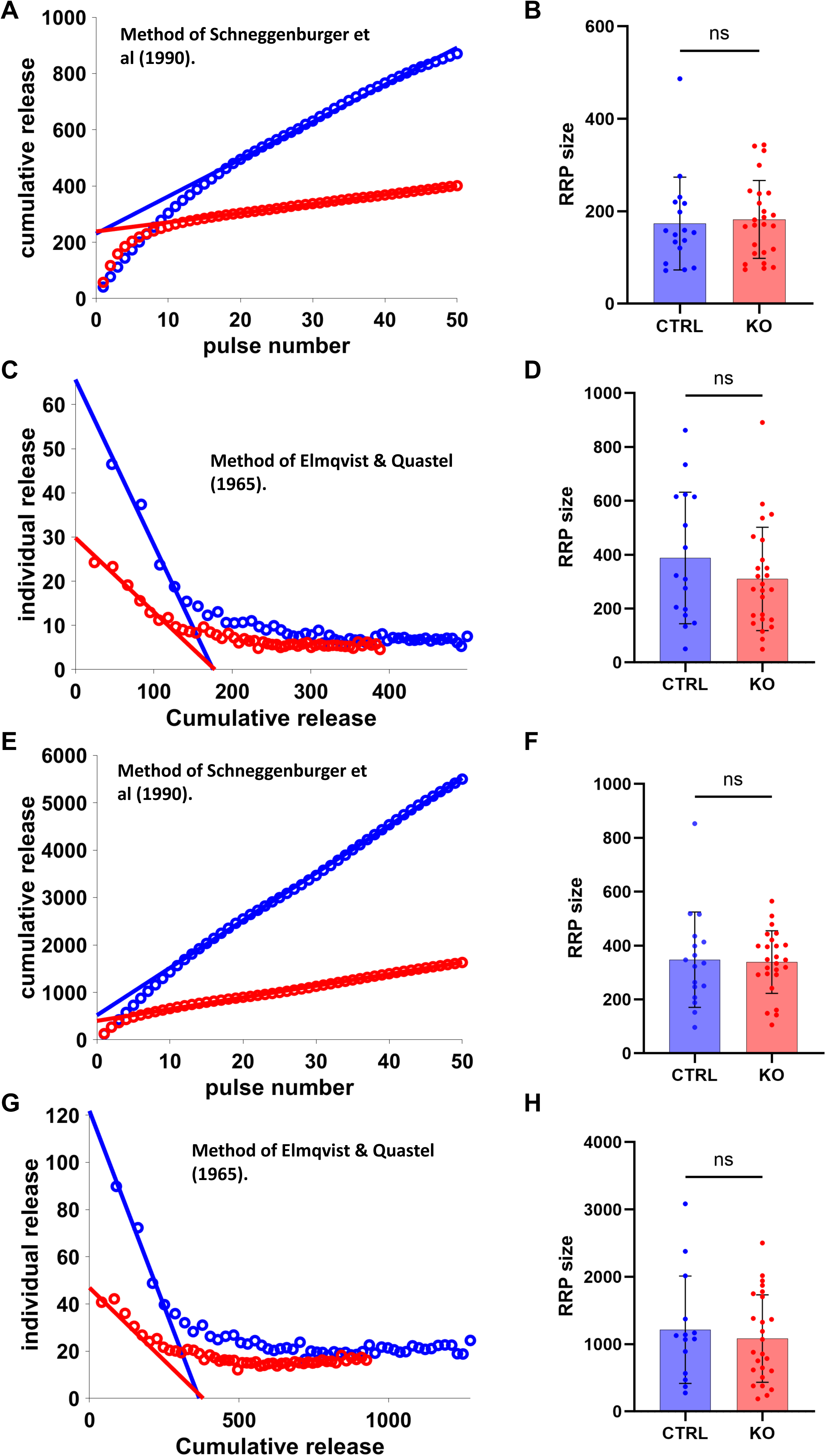
MCU deficiency does not alter RRP size at the endbulb of Held synapse. **A**. Example cumulative release plots from control and MCU KO mice using 400 Hz stimulus trains. Lines: linear fit to the steady-state release of the trains (15^th^ to 50^th^ pulses). RRP size was determined as the vesicle release at y-intercept. **B**. Summary of RRP sizes calculated using the method from Schneggenburger et al. (1999) from control and MCU KO mice. ns, not significant. **C**. Individual release vs cumulative release plot using 400 Hz stimulus train. Lines: linear fit of the early release events of the stimulus train. **D**. Summary of RRP sizes calculated using the method from Elmqvist and Quastel (1965) from control and MCU KO mice. ns, not significant. **E-F**: RRP size calculated based on eEPSC charge transfer of 400 Hz trains, using the method by Schneggenburger et al. (1999). **G-H**: RRP size calculated based on eEPSC charge transfer of 400 Hz trains, using the method by Elmqvist and Quastel (1965).

### MCU deficiency promotes synaptic recovery

To investigate the effect of MCU deficiency on synaptic recovery, we additionally recorded a post-train eEPSC at various delays (10-1500 ms) after the end of the 50-pulse train at 100 Hz (Fig. 5A, C) or 400 Hz (Fig. 5B, D). Recovery of eEPSC peak amplitude after 100 Hz stimulus train was not significantly different between control and MCU KO mice (Fig. 5E; Two-way ANOVA, Control, n = 21; MCU KO, n = 26; mice strain effect, *F_(_*_1,585)_ = 0.4415, P=0.5067; post-train delay effect: *F_(_*_12,585)_ = 7.603, P*<*0.0001; Interaction, *F_(_*_12,585)_ = 1.713, P = 0.0604). However, the recovery of eEPSC peak amplitude after 400 Hz trains were significantly different between two groups (Fig. 5F; two-way ANOVA, mice strain effect, *F_(_*_1,572)_ = 4.106, P = 0.0432; post-train delay effect, *F_(_*_12,572)_ = 25.68, P *<* 0.0001; Interaction, *F_(_*_12,572)_ = 3.365, P *<* 0.0001). Specifically, eEPSC peak amplitude recovered slower within the first 70 ms following the train in MCU KO mice (Fig. 5F), which suggests that it takes longer for synchronously released vesicles to recover at MCU deficient synapse. Consistently, we observed increased baseline response following the trains in MCU KO mice due to sustained asynchronous release (Fig. 5D).

**Figure 5:**
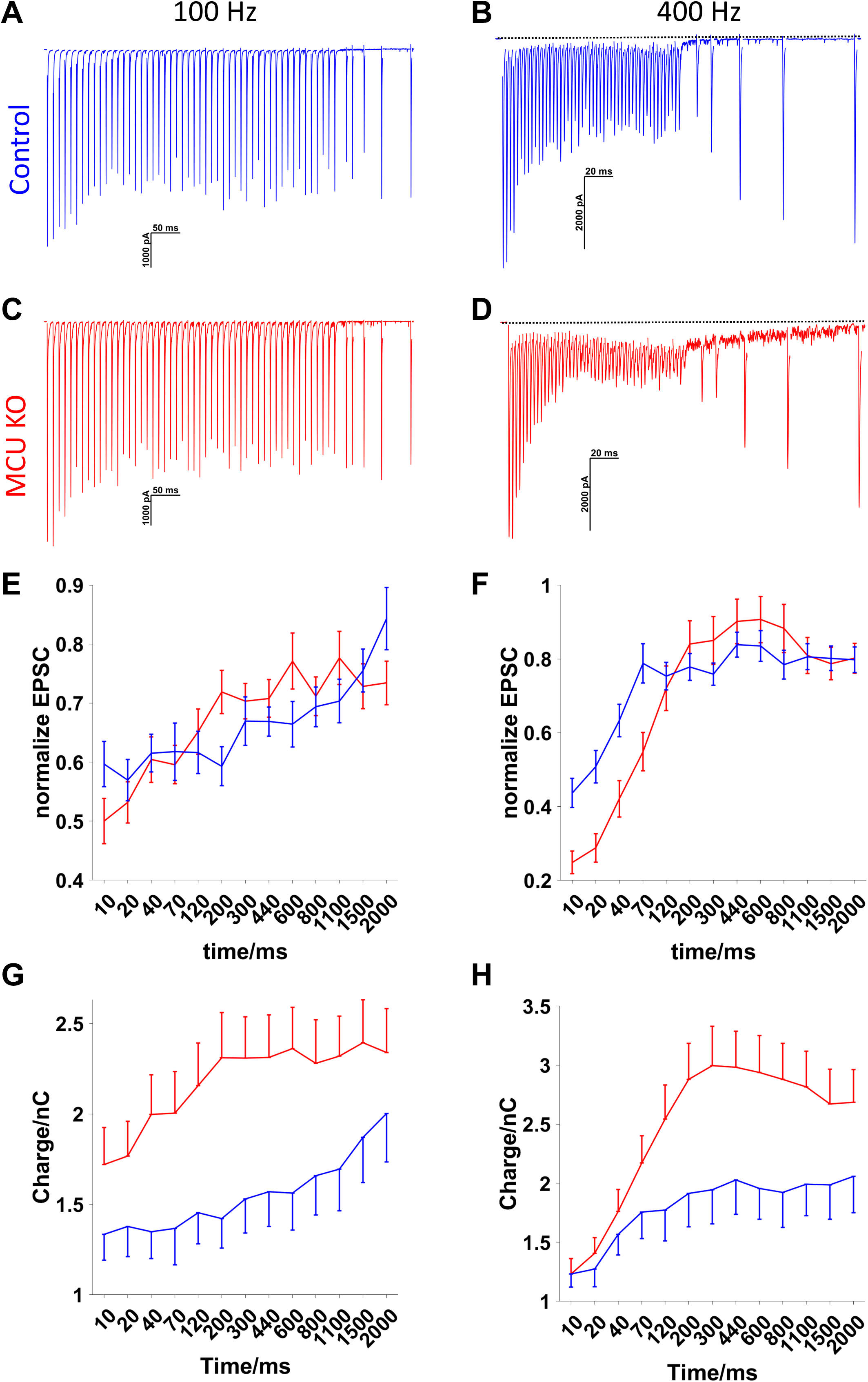
MCU deficiency promotes synaptic recovery. **A-D**, Example eEPSC recovery following 50 pulse stimulus trains at 100 Hz (**A, C**) and 400 Hz (**B, D**). **E-F**: Recovery of eEPSC peak amplitude after 50 pulse trains at 100 Hz (**E**) and 400 Hz (**F**). G-H: Recovery of eEPSC charge transfer after 50 pulse trains at 100 Hz (**G**) and 400 Hz (**H**).

To account for the total vesicle release, we further calculated synaptic recovery based on eEPSC charge transfer. In contrast to eEPSC peak amplitude (Fig. 5E-F), post-train eEPSC charge transfer was significantly higher in MCU KO mice following trains at both 100 Hz (Fig. 5G; Control: n = 21, MCU KO: n = 26; two-way ANOVA, mice strain effect: *F_(_*_1,585)_ = 52.07, P *<* 0.0001; post-train delay effect: *F_(_*_12,585)_ = 1.683, P = 0.0667; Interaction: *F_(_*_12,585)_ = 0.3018, P = 0.9891) and 400 Hz (Fig. 5H; Control: n = 20, MCU KO: n = 26; two-way ANOVA, mice strain effect: *F_(_*_1,572)_ = 39, P *<* 0.0001; post-train delay effect: *F_(_*_12,572)_ = 5.417, P *<* 0.0001; interaction: *F_(_*_12,572)_ = 0.8977, P = 0.5492). Overall, these results suggest that synaptic recovery in terms of total vesicle release is faster in MCU KO mice, presumably due to higher calcium accumulation at the endbulb of Held in these mice during sustained high-rate activity (Wang and Manis, 2008).

### MCU deficiency does not alter intrinsic membrane properties of post-synaptic bushy neurons

To further elucidate the function of MCU in auditory processing at this important neural connection, we next investigated the intrinsic membrane properties of the postsynaptic bushy neurons from control and MCU KO mice. Current-clamp recording was performed from target bushy neurons to acquire membrane responses to 200 ms current step injections at levels varied from-500 nA to 1000 nA in 25 nA increment (Fig. 6A). Resting membrane potential showed no significant difference between bushy neurons from control and MCU KO mice (Fig. 6B; Control, *−*64.54 *±* 1.725 mV, n = 17; MCU KO, *−*64.63 *±* 2.178 mV, n = 21; Mann-Whitney test, P = 0.7496). Membrane input resistance was calculated as the slope of the current-voltage plot of hyperpolarized responses to small current injections (from-25 to-100 pA), which showed no difference between two groups (Fig. 6C; Control, 50.62 *±* 19, 97 MΩ, n = 17; MCU KO, 51.69 *±* 13.72 MΩ, n = 21; Mann-Whitney test, P = 0.2909). These neurons also had similar membrane time constant (Fig. 6D; Control, 1.608 *±* 0.4561 ms, n = 17; MCU KO, 1.823 *±* 0.4876 ms, n = 21; Welch’s t test, P = 0.1696). We defined the threshold current of each neuron as the minimum current injection level required to trigger an action potential, which was also similar between control and MCU KO mice (Fig. 6E; Control, 369.1 *±* 179.3 nA, n = 17; MCU KO, 272.1 *±* 137 nA, n = 21; Mann-Whitney test, P = 0.1049). These results suggest that MCU deficiency does not alter intrinsic membrane properties of post-synaptic bushy neurons in the CN.

**Figure 6:**
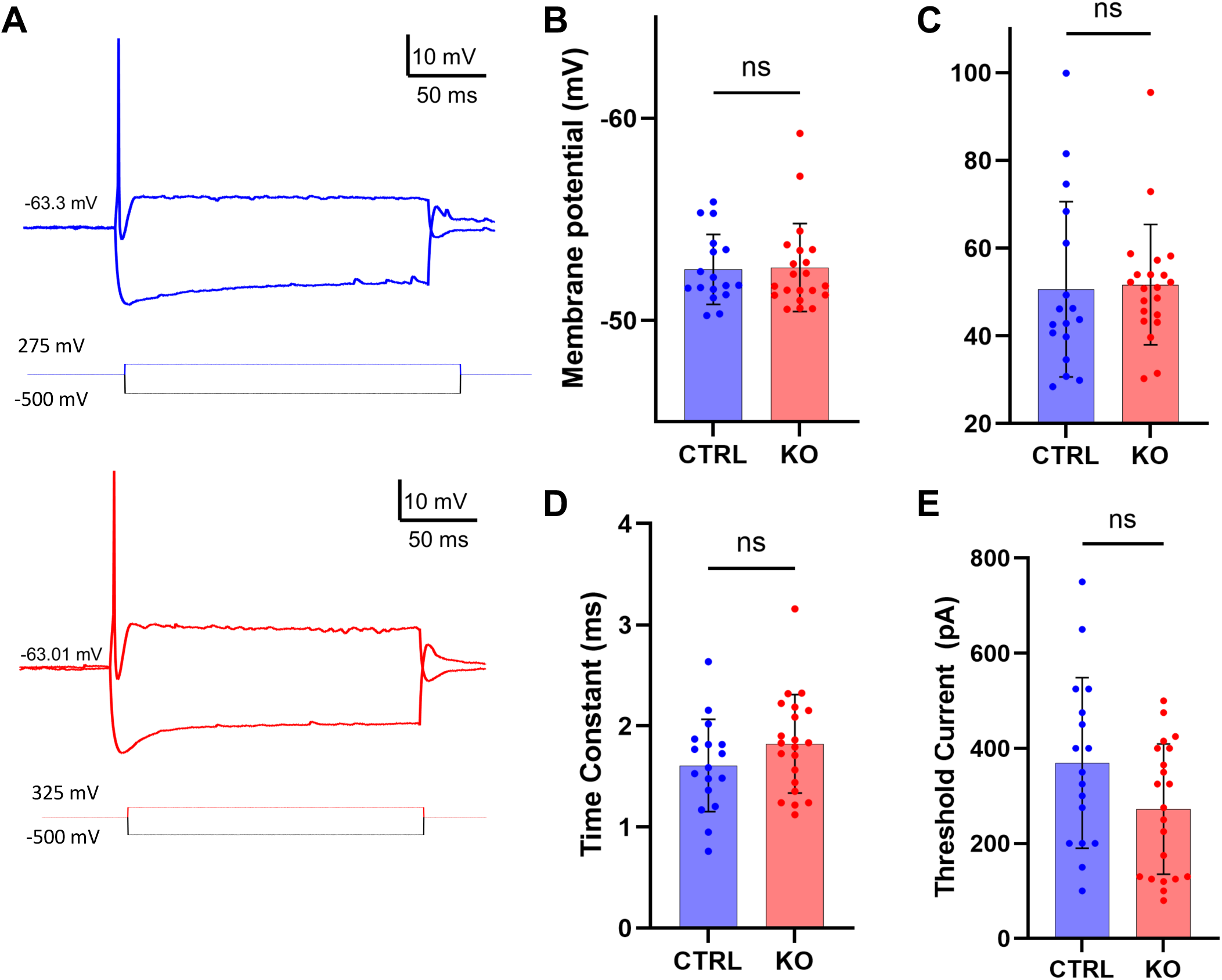
MCU deficiency does not alter intrinsic membrane properties of postsynaptic bushy neurons. **A**. Current injection evoked responses in example bushy neurons from control and MCU KO mice. Resting membrane potential was labeled in each neuron. **B-E**. Summary of all bushy neurons in resting membrane potential (**B**), input resistance (**C**), membrane time constant (**D**), and threshold current (**E**). ns, not significant.

### Temporal precision of spikes during high-rate activity is compromised in MCU KO mice

To investigate how MCU deficiency affects signal processing in bushy neurons, we examined the firing rate and spike timing of AN stimulation evoked spike trains at 100 and 400 Hz in control and MCU KO mice. We first recorded responses to 50-pulse AN stimulus trains at 100 and 400 Hz (Fig. 7A, B). At 100 Hz, AN stimulation evoked reliable spike trains in most neurons in both groups (Fig. 7A), with a slightly higher firing rate in MCU KO mice (Fig. 7C; Control, 85.85%*±*23.92%, n = 17; MCU KO, 92.24%*±*22.19%, n = 21; Mann-Whitney test, P = 0.0314). Temporal precision of spikes was evaluated by calculating the vector strength of the spike trains (Goldberg and Brown, 1969; Xie, 2016), which was not different between bushy neurons from control and MCU KO mice (Fig. 7D; Control, 0.9983 *±* 0.0021, n = 17; MCU KO, 0.9985 *±* 0.0024, n = 21; Mann-Whitney test, P = 0.8788). At 400 Hz, AN stimulation evoked spikes with lots of failures in both groups (Fig. 7B). Despite the high variation, the firing rate of spike trains was not statistically different between control and MCU KO (Fig. 7E; Control, 39.31% *±* 37.98%, n = 17; MCU KO, 59.63% *±* 30.85%, n = 19; Welch’s t test, P = 0.0902). In contrast, vector strength of spikes was significantly reduced in MCU KO mice (Fig. 7F; Control, 0.9499*±*0.0551, n = 17; MCU KO, 0.8766*±*0.1186, n = 19; Mann-Whitney test, P = 0.0194). These results likely reflect substantial variability in the dynamic balance between synchronous and asynchronous vesicle releases during the short 50-pulse stimulus trains at different rates (Figs. 3, 5).

**Figure 7:**
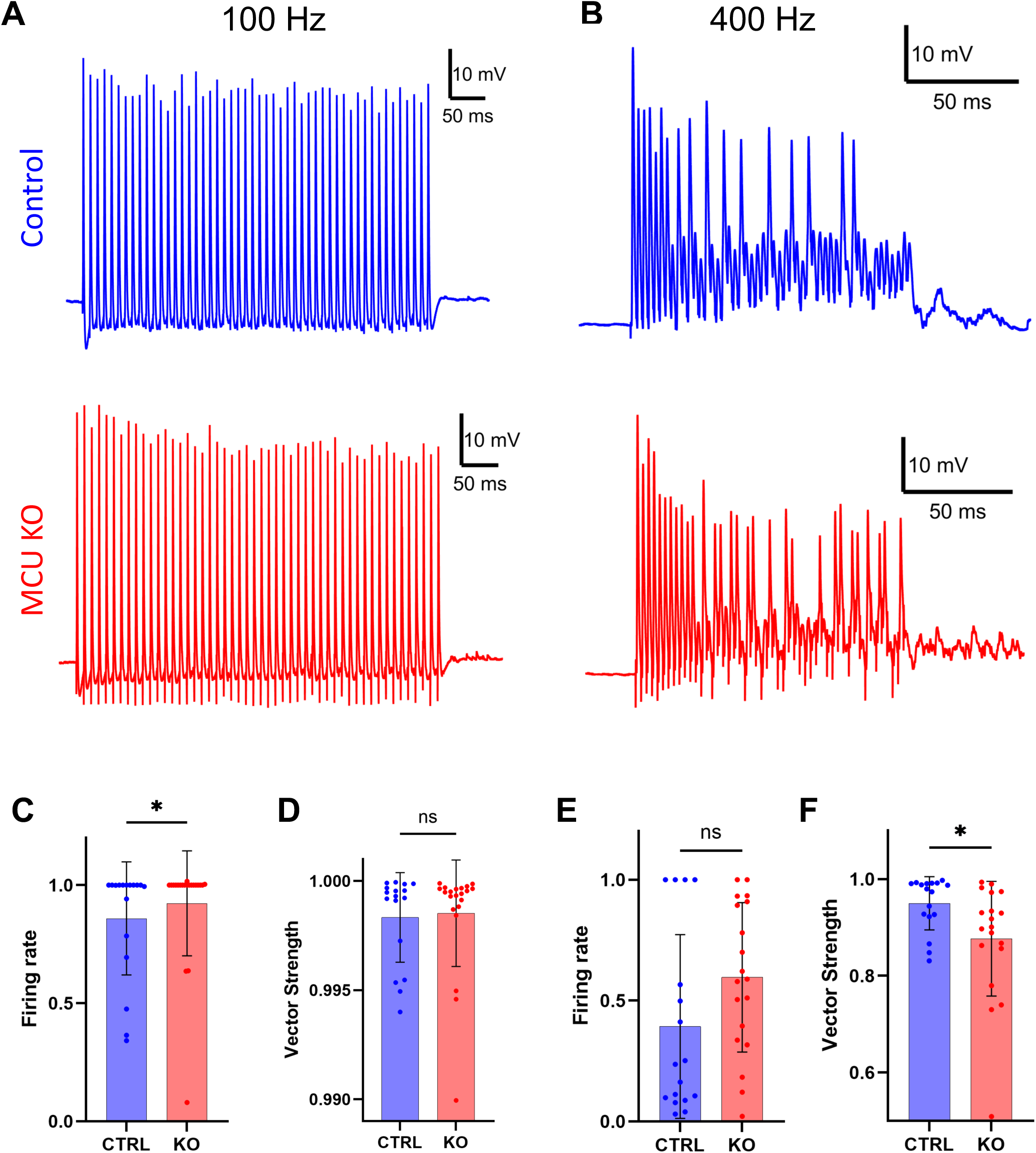
-MCU deficiency affects firing rate and spike timing of 50 pulse trains. **A-B**: Spike trains evoked by AN stimulation at 100 Hz (**A**) and 400 Hz (**B**) in example bushy neurons from control and MCU KO mice. **C-D**: Summary data in firing rate and vector strength from 100 Hz spike trains. **E-F**: summary data in firing rate and vector strength from 400 Hz spike trains. ns, not significant; *P*<*0.05

Auditory neurons are known to sustain high-rate activity over prolonged time under physiological condition. In order to evaluate the effect of MCU deficiency under those activity levels, we further assessed spikes trains in bushy neurons evoked by 900-pulse AN stimulus trains at 100 and 400 Hz (Fig. 8). Bushy neurons from both control and MCU KO mice were able to fire reliable spikes throughout the long trains at 100 Hz (Fig. 8A), but showed high failure rates at 400 Hz (Fig. 8B). Interestingly, the overall firing rate was not significantly different between control and MCU KO at either 100 Hz (Fig. 8C; Control, 76.62% *±* 24.9%, n = 14; MCU KO, 68.48% *±* 31.96%, n = 17; Welch’s t test, P = 0.4313) or 400 Hz (Fig. 8E; Control, 20.33% *±* 28.47%, n = 12; MCU KO, 10.53% *±* 7.44%, n = 16; Welch’s t test, P = 0.2670). In contrast, the vector strength of evoked spikes was significantly reduced in MCU KO mice in prolonged trains at both 100 Hz (Fig. 8D; Control, 0.9976 *±* 0.0024, n = 14; MCU KO, 0.9934 *±* 0.0068, n = 17; Mann-Whitney test, P = 0.0078) and 400 Hz (Fig. 8F; Control, 0.8750 *±* 0.0560, n = 12; MCU KO, 0.6876 *±* 0.2148, n = 16; Welch’s t test, P = 0.0036). The results suggest that during prolonged high-rate activity, loss of MCU does not affect the firing rate but decreases the temporal precision of AN evoked spikes in bushy neurons. Overall, we conclude that MCU functions to regulate calcium at the endbulb of Held synapse and dynamically modulate synaptic vesicle release during sustained high-rate activity, subsequently improves auditory temporal processing.

**Figure 8:**
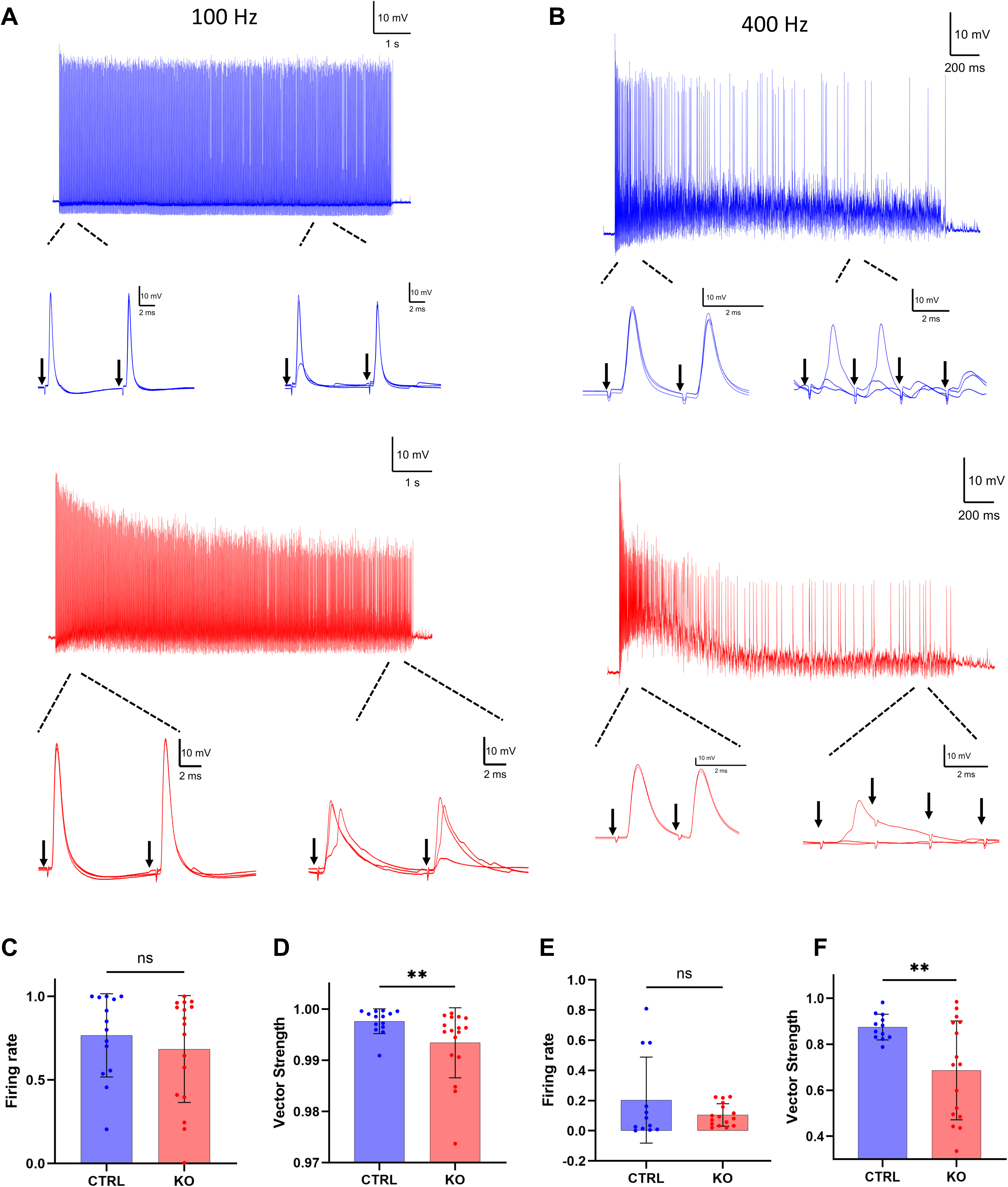
Temporal precision of spikes during prolonged high-rate activity is compromised in MCU KO mice. **A-B**: Prolonged spike trains evoked by 900 pulse stimulation at 100 Hz (**A**) and 400 Hz (**B**) in example bushy neurons from control and MCU KO mice. Insets show magnified view of spikes at the beginning and end of the trains. **C-D**: Summary data in firing rate and vector strength from prolonged 100 Hz trains. **E-F**: summary data in firing rate and vector strength from prolonged 400 Hz trains. ns, not significant; **P*<*0.01.

## DISCUSSION

Calcium accumulation at the synaptic terminal is rate dependent, with higher activity leading to faster and greater increases in intracellular calcium concentration (Neher and Sakaba, 2008). Given that MCU is activated only above a certain calcium threshold, its function is likely tightly coupled to neural activity. Previous studies investigating MCU function either studied cultured neurons (Marland et al., 2016; Ashrafi et al., 2020) or investigated only local field potentials (Kim et al., 2011; Devine et al., 2022; Pannoni et al., 2025). The roles of MCU in synaptic transmission and neural processing under physiologically relevant activity levels remain unclear. In this study, we demonstrated rate-dependent function of MCU at the endbulb of Held synapse and postsynaptic bushy neuron connection. Our findings showed that MCU deficiency did not change the basal synaptic transmission of the endbulb or the intrinsic membrane properties of the postsynaptic bushy neurons. During sustained high-rate activity, however, synaptic transmission and functional output of the connection were significantly altered in MCU KO mice. In particular, we showed that MCU deficiency impairs auditory temporal processing, which may contribute to central auditory processing deficits and ultimately hearing loss.

### Function of MCU in synaptic transmission

Synaptic transmission starts from presynaptic action potential invasion, which activates voltage-gated calcium channels and generate a brief hike of calcium in the immediate vicinity of the channels that triggers vesicle exocytosis (Neher and Sakaba, 2008). Free calcium ions are quickly removed to the basal level and exocytosis stops. During sustained activities, repetitive calcium inflex lead to accumulation of free calcium ions that spread to larger domains in the synaptic terminal and affect the process of synaptic transmission (Zucker and Regehr, 2002). Under normal conditions, calcium accumulation is effectively cleared via multiple pathways, including passive diffusion and buffering, uptake into the endoplasmic reticulum by sarcoendoplasmic reticulum Ca^2+^-ATPase (SERCA) (Kim et al., 2005), extrusion across the plasma membrane by plasma membrane CA2+-ATPase (PMCA) (Zenisek and Matthews, 2000; Kim et al., 2005), removal by sodium/calcium exchanger (NCX) to extracellular space (Reuter and Porzig, 1995), and particularly MCU uptake into mitochondria (Billups and Forsythe, 2002; Marchi and Pinton, 2014), which was reported to account for up to 40% of presynaptic calcium clearance (Kim et al., 2005; Shutov et al., 2013). As a low-affinity, high-capacity calcium buffer, MCU is not activated at resting calcium level (∼ 100 nM) (Gleichmann and Mattson, 2011; Garbincius et al., 2020), but only uptakes calcium into mitochondria when cytosolic calcium reaches a high concentration (Kim et al., 2005; Marchi and Pinton, 2014; Finkel et al., 2015) (also see (Ashrafi et al., 2020)). Indeed, our findings showed that MCU functions in an activity rate-dependent manner in that MCU deficiency did not affect basal synaptic properties including quantal size, quantal content, release probability (Fig. 2), as well as RRP size (Fig. 4). During sustained high-rate activity, we found that synaptic transmission was significantly shifted toward increased asynchronous release as activity progressed (Fig. 3G-H). Under such conditions, synaptic calcium accumulates to excessively high levels in the microdomain of the synaptic terminal, resulting in enhanced asynchronous release of synaptic vesicles (Kaeser and Regehr, 2014). In the absence of MCU-mediated uptake, elevated synaptic calcium in MCU KO mice also leads to increased overall vesicle release at the beginning of the eEPSC train (Fig. 3E-F), but this effect quickly diminishes presumably due to accelerated depletion of the RRP.

### Role of MCU in shaping the functional output of postsynaptic bushy neurons

Effective signal transmission depends not only on presynaptic vesicle release but also on the generation of membrane responses in postsynaptic neurons. Despite the abundance of mitochondria in bushy neurons (Spirou et al., 2023), we did not observe any significant differences in their intrinsic membrane properties between control and MCU KO mice (Fig. 6), indicating that MCU deficiency does not alter postsynaptic excitability under resting condition. Unexpectedly, when we used 50-pulse AN stimulus trains, we found that the firing rate of 100 Hz trains was increased in MCU KO mice without significant changes in spike timing (Fig. 7C-D), whereas the firing rate of 400 Hz trains was not changed but spike timing was significantly compromised (Fig. 7E-F). The seemingly complex results likely reflect the dynamic interplay among rate-dependent calcium accumulation, enhanced vesicle release, accelerated depletion and replenishment of RRP, and shifts in the relative contributions of synchronous and asynchronous vesicle release. To minimize the influence of these transient synaptic dynamics and better assess sustained transmission, we further examined the spike output of bushy neurons with prolonged 900-pulse AN stimulus trains, a physiologically relevant paradigm given the continuous activity of the auditory system. Indeed, the results showed that MCU deficiency had no significant effect on prolonged spike trains at either 100 or 400 Hz, but significantly reduced the temporal precision of evoked spikes (Fig. 8C-F). Overall, the results showed that MCU functions to promote auditory temporal processing at this endbulb of Held / bushy neuron connection.

### MCU and hearing loss

The auditory system relies on the temporal information of sound to sense the acoustic environment, including performing tasks like sound localization (Joris and Yin, 2007) and pitch detections (Shofner, 2008). Hearing loss is widely associated with impaired temporal processing in the central auditory nervous system (Lorenzi et al., 2006; Lorenzi et al., 2009; Grose and Mamo, 2010; Anderson et al., 2012), a key component of central auditory processing deficits (Atcherson et al., 2015). In particular, bushy neurons are among the major cell types that encode information about temporal fine structure of sound (Joris et al., 1994b; Joris et al., 1994a), which provide temporal cues upward to support the auditory function. Impaired signal processing in bushy neurons has been reported in various pathological conditions associated with hearing loss (Wang and Manis, 2006; Manis et al., 2011; Garcia-Hernandez et al., 2017; Xie et al., 2024; Ryugo and Nishitani, 2026). Our findings in this study that bushy neurons in MCU KO mice show similar firing rates but reduced temporal precision during sustained high-rate activity suggest that compromised MCU function may be an important mechanism that contributes to central auditory processing deficits, and ultimately to the development of hearing loss.

## Author Contributions

G.L. and R.X. designed research; G.L. performed research and analyzed data; G.L. and R.X. wrote the paper.

## Acknowledgements

This work was supported by NIH grants R01DC020582 and P01AG051443. We thank Dr. Shengyin Lin for his general lab support and animal management.

